# Identification and Optimization of Kratom Strictosidine Pathway Enabled by Yeast Multiplex Engineering

**DOI:** 10.64898/2026.04.16.719034

**Authors:** Yinan Wu, Dong Oh Han, Franklin Leyang Gong, Sijin Li

**Author notes:** These authors contributed equally (Yinan Wu, Dong Oh Han). Corresponding Author: Sijin Li.

## Abstract

Monoterpene indole alkaloids (MIAs) are a major class of plant natural products with important pharmaceutical activities, yet the biosynthetic pathway to their universal precursor, strictosidine, has been fully elucidated in only Catharanthus roseus. In kratom (Mitragyna speciosa), only the first and last steps of strictosidine biosynthesis were previously known. Here, we applied multiplex pathway engineering in yeast to accelerate the discovery, reconstruction, and optimization of the kratom strictosidine pathway. Iterative multiplex integration and screening identified 13 functional kratom genes and enabled rapid validation of functional pathway modules, thereby completing the kratom strictosidine pathway from geranyl pyrophosphate and tryptophan. We also identified a vacuolar secologanin transporter, MsNPF2.6, which increased strictosidine production by 62% in yeast. Pathway optimization through the incorporation of nepetalactol-producing enzymes from other plants further supported strictosidine production in yeast from fed geraniol and tryptophan. These results establish the strictosidine pathway in kratom and highlight multiplex engineering as a powerful platform for rapid plant pathway discovery and optimization.

## Introduction

Monoterpene indole alkaloids (MIAs) represent one of the largest and most structurally diverse classes of plant natural products (PNPs) with great pharmaceutical potential^1,2^. Examples include the anticancer agents vincristine and vinblastine, the antimalarial quinine, and the antihypertensive reserpine. To date, more than 3,000 MIAs have been identified from plants, primarily in the order Gentianales, especially within the families Apocynaceae, Rubiaceae, and Loganiaceae^3–8^. Most MIAs derive from the universal precursor, strictosidine, the biosynthetic mechanism of which has only been fully elucidated in the model MIA-producer, *Catharanthus roseus*, in the Apocynaceae family^3,9^. This pathway contains 13 steps and two metabolic branches that convert tryptophan to tryptamine and geranyl pyrophosphate (GPP) to secologanin, respectively, which are finally condensed by strictosidine synthase (STR). Emerging studies have focused on elucidating pathways downstream of strictosidine that lead to novel MIAs, including mitragynine^6,10,11^, mitraphylline^12^, ibogaine^13,14^, and voacangine^15,16^. In the meantime, the understanding of the strictosidine biosynthetic pathway has remained limited. Enzymes catalyzing individual reactions have been reported in a few MIA-producing plants, including STRs from *Rauvolfia serpentina*^17^, *Ophiorrhiza pumila*^18^, *Camptotheca acuminata*^19^, as well as enzymes involved in iridoid synthesis from *Vinca minor*^20,21^, Nepeta (catmint)^22–24^, and *Olea europaea* (olive)^25^. Yet *C. roseus* remains the only species with an experimentally validated strictosidine pathway and the foundation of all subsequent MIA studies.

Kratom (*Mitragyna speciosa*), a member of the Rubiaceae family, has recently emerged as another important MIA producer^6,10,26–28^. It has independently evolved to produce a distinct repertoire of alkaloids, most prominently the analgesic mitragynine and its stereoisomer speciogynine. Kratom also exhibits a prominent 3R epimerization machinery that contributes to the synthesis of numerous spirooxindole alkaloids^29,30^. These MIAs highlight the plasticity of MIA biosynthesis in kratom and its role as a comparative example for investigating the conserveness and diversification of MIA biosynthesis among various plants. To date, only two enzymes that catalyze the first and last steps in strictosidine synthesis, including MsTDC (tryptophan decarboxylase) and MsSTR, have been fully identified and characterized from kratom^11,31,32^. All other enzymes remain unknown despite the availability of sequenced transcriptomes and genomes^10,11,33,34^, as well as the well-characterized *C. roseus* pathway, which provides the biochemical framework. This gap primarily results from the time-consuming and laborious process of elucidating pathways in non-model, medicinal plants. Given the challenges in plant engineering, heterologous approaches to characterize a putative PNP pathway in a versatile host such as yeast^35,36^ or *Nicotiana benthamiana*^30,37,38^ have proven highly effective. However, these approaches are typically pursued in a stepwise, gene-by-gene manner because the characterization of one enzyme requires a sufficient supply of its precursor *in vivo*, achieved by characterizing and expressing the upstream enzymes. As a typical strictosidine biosynthetic pathway likely involves 11 enzymatic reactions, along with auxiliary proteins such as transporters and redox partners^9,21,39–43^, the stepwise characterization strategy is inherently slow and laborious.

In this study, we leveraged a previously developed multiplex gene assembly method^44^ to accelerate the elucidation, reconstruction, and optimization of complex PNP pathways, using the kratom strictosidine pathway as an example (**Fig. 1**). The first two rounds of multiplex gene assembly validated the function of 10 enzymes catalyzing the conversion from endogenous geranyl pyrophosphate (GPP) to secologanin, including a geraniol synthase (GES), a geraniol-8-hydroxylases (G8H), an iridoid oxidase (IO), a 7-deoxyloganetic acid glucosyltransferase (7DLGT), a 7-deoxyloganic acid hydroxylase (7DLH), two loganic acid methyltransferases (LAMT), a secologanin synthase (SLS), a cytochrome P450 reductase (CPR), and a cytochrome P450-associated alcohol dehydrogenase (CYPADH), leaving only the enzymes converting 8-hydrogeraniol to nepetalactone unresolved. Following the recent report of an iridoid cyclase (ICYC) in multiple plants^45^, we then identified and functionally characterized an iridoid synthase (ISY) and an ICYC in kratom. Along with the discovery of an 8-hydroxygeraniol oxidoreductase (8HGO)-like enzyme identified in a separate study, we have found at least one functional enzyme at each step of the kratom biosynthetic pathway.

**Figure 1.**
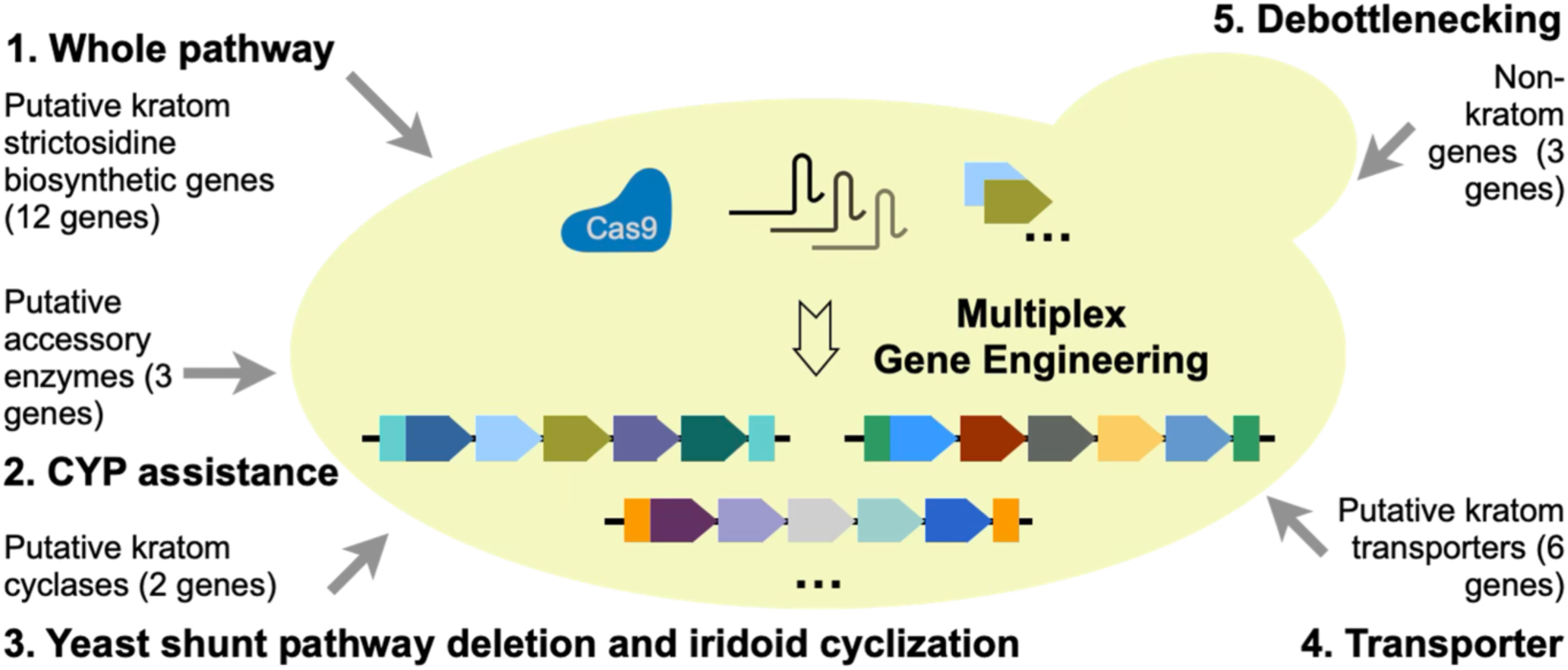
Schematic of identification and optimization of kratom strictosidine pathway in yeast with multiplex engineering. Candidate genes encoding putative kratom strictosidine biosynthetic enzymes, CYP assistant enzymes, iridoid cyclases, and transporters, were expressed and functionally identified in yeast with the MULTI-SCULPT multiplex engineering approach that enabled integration of up to 12 genes per round. Non-kratom genes were also integrated to debottleneck the rate-limiting steps.

We leveraged the multiplex method to identify transporters involved in strictosidine synthesis and discovered a novel vacuolar secologanin transporter, MsNPF2.6, which enhanced strictosidine production by 62%. We finally engineered the yeast to produce strictosidine from geraniol and tryptophan by complementing the kratom pathway with three active, non-kratom genes from catnip and *V. minor*, including Vmi8HGO-A, NcaISY, and NcaMLPLA, which encodes a nepeta-specific iridoid cyclase^21,24^. In summary, this work discovered and characterized 13 functional kratom genes and one transporter involved in strictosidine biosynthesis by combining transcriptomics with large-scale multiplex pathway integration in yeast. This integration-based strategy provides a rapid characterization method that complements existing transcriptomics-driven gene prediction approaches to accelerate PNP pathway discovery.

## Results

### Elucidating early and late steps of the kratom strictosidine pathway in yeast using full pathway assembly

The elucidated strictosidine biosynthetic pathway in *C. roseus* involves 12 enzymes that collectively catalyze the conversion of geranyl pyrophosphate and tryptophan into strictosidine (**Fig. 2a** and **2b**)^9,39^. While their enzymatic functions have been extensively studied in *C. roseus*, most remain uncharacterized in kratom, except for MsTDC and MsSTR^11,31,32^. Using the recently published kratom transcriptome^11,33^, we mined for homologs of the nine remaining enzymes based on *C. roseus* enzyme sequences (**Table S1-S3**), including GES, G8H, 8HGO (also called 10HGO or GOR in prior studies), ISY, IO, 7DLGT, 7DLH, LAMT, and SLS. Single candidates exhibiting high sequence identity (>75%) were identified for six out of the nine enzymes (GES, ISY, IO, 7DLGT, 7DLH, and SLS), and were therefore designated MsGES, MsISY, MsIO, Ms7DLGT, Ms7DLH, and MsSLS. Two isoforms with high sequence identity (87%) were identified for 8HGO, originating from alternative start codons within the same transcript. Furthermore, two distinct candidates were identified for G8H and LAMT: MsG8H1 and MsG8H2 (82% and 74% identity, respectively), and MsLAMT1 and MsLAMT2 (79% and 81% identity, respectively). Most of these candidates showed higher expression in young leaves, consistent with the higher accumulation of strictosidine in these tissues (**Fig. S1**)^11^.

**Figure 2.**
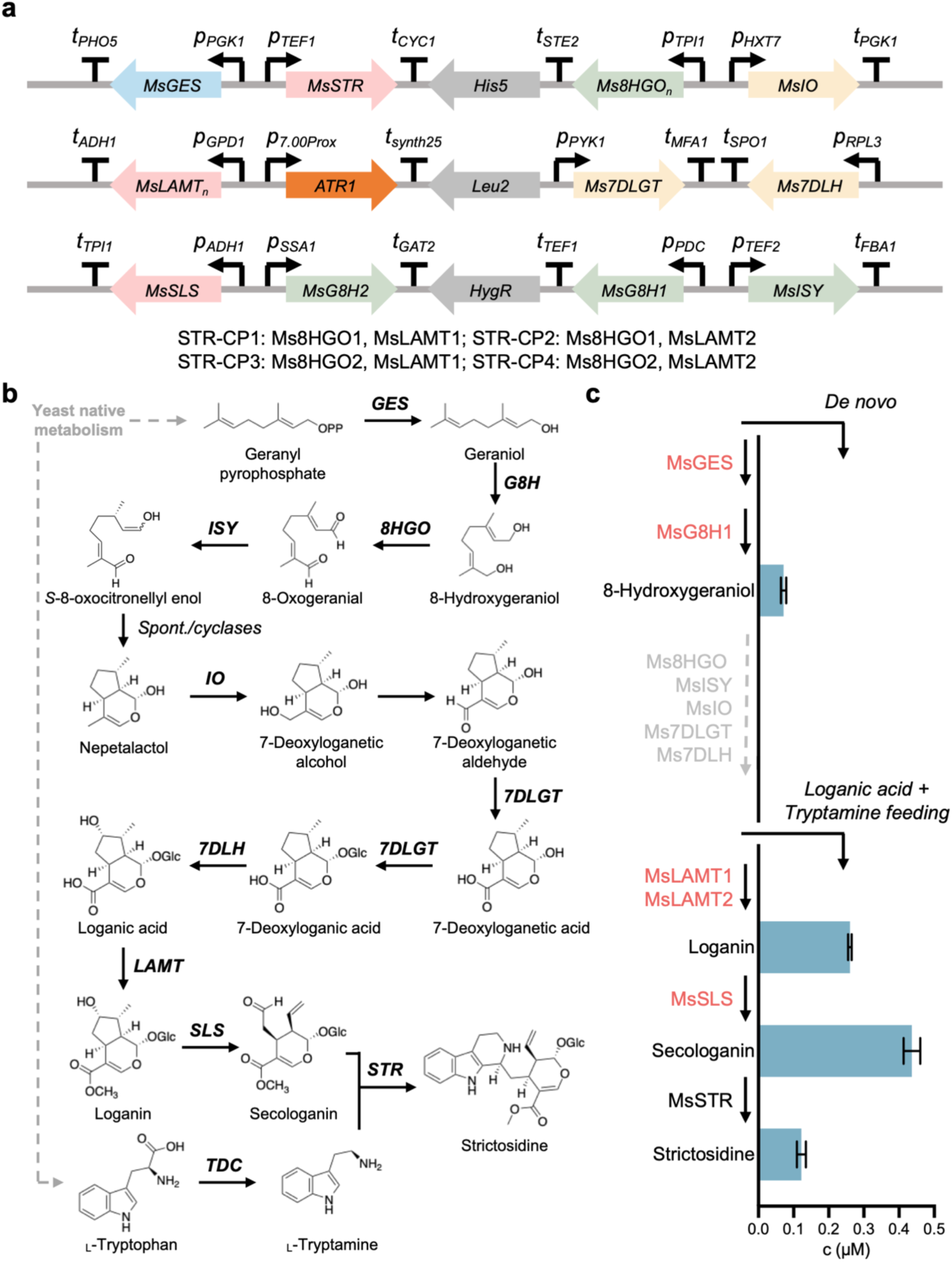
Validation of kratom strictosidine biosynthetic enzymes at early and late steps after whole pathway assembly with multiplex engineering. **a.** Whole pathway assembly in yeast using the MULTI-SCULPT multiplex engineering methods for characterization of strictosidine biosynthetic candidate enzymes, resulting STR-CP1, STR-CP2, STR-CP3, and STR-CP4 strains. **b.** The strictosidine biosynthetic pathways in yeast. c. *De novo* and loganic acid and tryptamine feeding assays for STR-CP3, validating the function of MsGES, MsG8H1, MsLAMT1, MsLAMT2, and MsSLS (highlighted in red). Data are presented as mean ± s.d. from three independent biological replicates.

The MULTI-SCULPT method we previously developed can integrate up to 12 biosynthetic genes into the yeast genome^44^. To rapidly identify functional enzymes among these candidates, we used the MULTI-SCULPT method to integrate the entire putative secologanin biosynthetic pathway into the yeast strain CEN.PK2-1D (referred to as CEN), together with the characterized MsSTR and a CPR from *Arabidopsis thaliana* (ATR1), and generated four engineered strains (STR-CP1, STR-CP2, STR-CP3, and STR-CP4) (**Fig. 2a**). Genes were expressed using specific promoter-terminator pairs with medium to strong expression strengths (**Fig. 2a** and **Fig. S2**). Since the conversion of geraniol to 8-hydroxygeraniol catalyzed by G8H has been identified as the rate-limiting step in strictosidine biosynthesis^39^, each strain contained both *MsG8H1* and *MsG8H2*, along with one copy of the remaining pathway genes, including different combinations of *Ms8HGO1/2* and *MsLAMT1/2*. *MsSTR* was included to enable strictosidine biosynthesis from secologanin and exogenously supplied tryptamine, while *ATR1* was incorporated to maintain redox balance and support the activity of cytochrome P450 (CYP) enzymes (G8H, IO, 7DLH, and SLS) in the pathway^46^.

To validate the predicted enzymes *in vivo*, we fed tryptamine to the STR-CP strains, using the parental CEN strain as a negative control. To account for the possibility that certain early-step enzymes were active but produced insufficient metabolic flux for downstream reactions, we also supplemented tryptamine in combination with geraniol, nepetalactol, or loganic acid as alternative starting precursors. After 72 hours of fermentation, culture supernatants were collected and analyzed by HPLC/Q-TOF mass spectrometry (MS) to quantify strictosidine and pathway intermediates (**Fig. S3**). The production of loganin, secologanin, and strictosidine from loganic acid and tryptamine across all STR-CP strains, but not the CEN control, validated the function of MsLAMT1, MsLAMT2, and MsSLS, which catalyze the conversion of loganic acid to secologanin (**Fig. 2c** and **Fig. S4**). *De novo* production of 8-hydroxygeraniol (∼0.07 μM) was observed in STR-CP3 and STR-CP4 strains, validating the function of MsGES and one or both of MsG8H1 and MsG8H2 (**Fig. 2c** and **Fig. S4**). When supplemented with geraniol, STR-CP3 and STR-CP4 produced markedly higher titers of 8-hydroxygeraniol (>2.42 μM), further confirming G8H activity (**Fig. S4**). However, no 8-oxogeranial or downstream intermediates were detected. Instead, MS peaks with *m/z* of 157.1587, 199.1693, and 175.1693 corresponding to yeast shunt pathway byproducts^47–49^ were observed (**Fig. S5**).

To further characterize MsG8H1 and MsG8H2, which were co-expressed in all STR-CP strains, we co-expressed *MsG8H1*, *MsG8H2*, or *CrG8H* together with *ATR1* in the CEN strain using yeast expression plasmids. Geraniol feeding assays revealed that MsG8H1 catalyzed the conversion of geraniol to 8-hydroxygeraniol, consistent with the activity of CrG8H, whereas MsG8H2 exhibited no detectable activity (**Fig. S6**).

Taken together, we identified five functional enzymes involved in the kratom strictosidine biosynthetic pathway at early and late steps (**Fig. 2c**, highlighted in red): MsGES, MsG8H1, MsLAMT1, MsLAMT2, and MsSLS. The activity of MsISY and Ms8HGO remain unclear.

### Validation of the middle step enzymes facilitated by the discovery of accessory enzymes

Feeding nepetalactol and tryptamine did not yield any downstream intermediates or strictosidine (Fig. S4), indicating a lack of activity in the middle steps of kratom strictosidine biosynthesis, which convert nepetalactol to loganic acid. These steps involve two CYPs, IO and 7DLH. Previous studies of the *C. roseus* pathway have shown that the accessory enzymes CrCPR, CrCYB5 (a cytochrome b5), and CrCYPADH (a CYP-associated alcohol dehydrogenase) are critical for these middle steps in yeast^21,39,50^. CrCPR and CrCYB5 enhance CYP activity, while CrCYPADH improves the spontaneous conversion of 7-deoxyloganetic alcohol to 7-deoxyloganetic acid in yeast.

Given the importance of accessory enzymes in loganic acid synthesis, we hypothesized that CYP accessory enzymes containing CPR, CYB5, and CYPADH would be necessary to provide additional redox partners and enable the functional validation of the middle steps. To discover the CYP accessory enzymes from kratom, we mined the kratom transcriptome using *C. roseus* protein sequences and identified one candidate each for CPR, CYB5, and CYPADH, which we named MsCPR, MsCYB5, and MsCYPADH (**Table S1-S3**). All three candidates showed medium to high expression across tissues (**Fig. S1**). We performed marker rescues on STR-CP3 to generate STR-CP5 and then integrated the accessory candidates, yielding the STR-MsCY strain. A positive-control strain, STR-CrCY, expressing the *C. roseus* accessory enzymes, was constructed in parallel.

To validate the function of both the middle-step and the accessory enzymes, we performed nepetalactol and tryptamine feeding assays. Although peaks corresponding to 7-deoxyloganetic alcohol, 7-deoxyloganetic aldehyde, and 7-deoxyloganetic acid were challenging to detect as reported previously, a novel putative peak corresponding to 7-deoxyloganic acid was observed in the fermentation products from both STR-CrCY and STR-MsCY strains (**Fig. 3a**). Additionally, both strains also produced loganin (∼0.06 μM), secologanin (∼0.28 μM), and strictosidine (∼0.05 μM). These results confirmed that MsIO, Ms7DLGT, and Ms7DLH are functional but require accessory enzymes to function in yeast.

**Figure 3.**
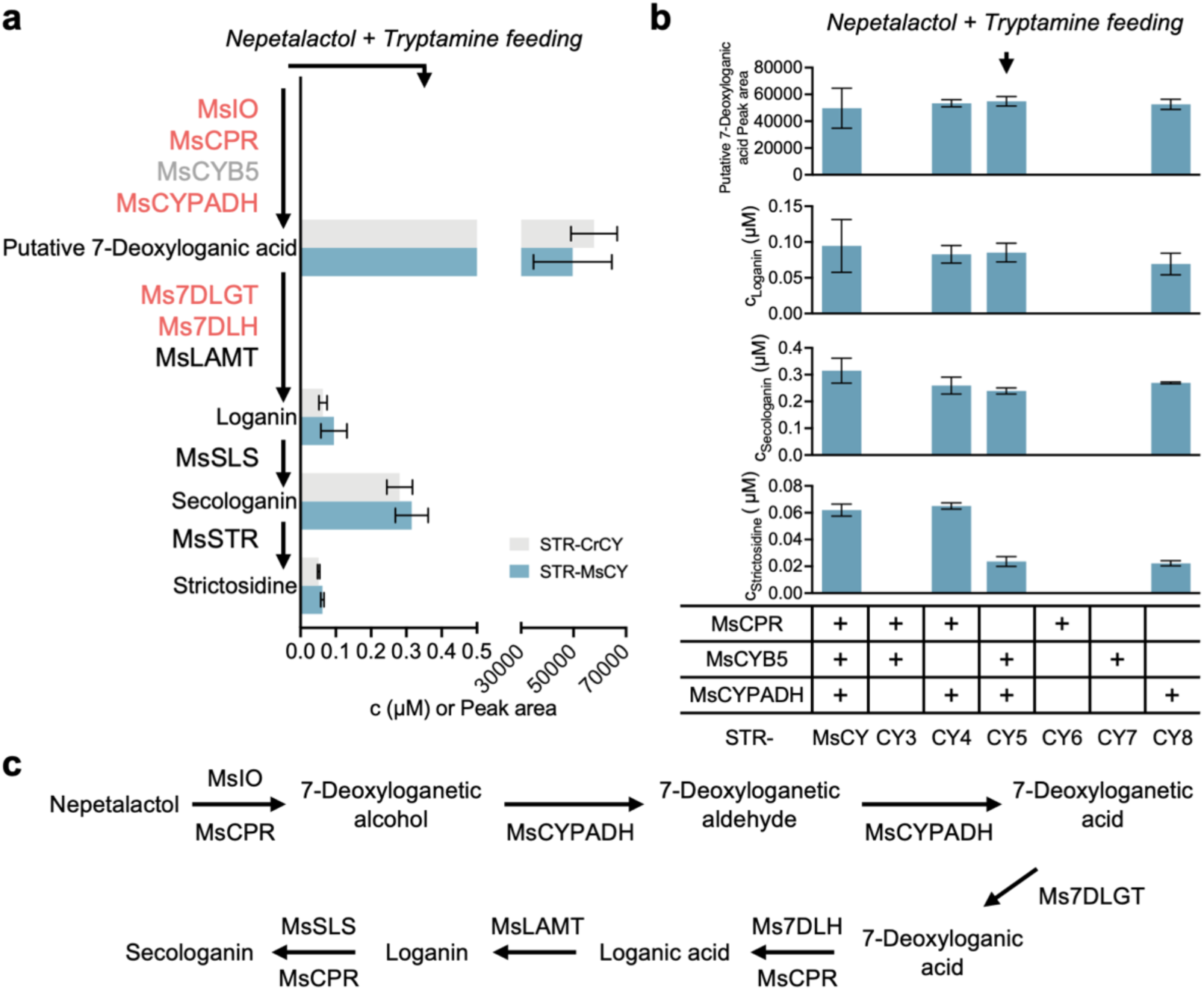
Validation of kratom strictosidine biosynthetic enzymes in middle steps together with CYP accessory enzymes. **a.** Results of nepetalactol and tryptamine feeding assays for STR-CrCY and STR-MsCY, each incorporating *C. roseus* and kratom CYP accessory enzymes (CPR, CYB5, and CYPADH) into the STR-CP3 strain following marker resuce. **b**. Results of nepetalactol and tryptamine feeding assays for strains harboring different combinations of kratom CYP accessory enzymes. **c.** Proposed functions of the five identified enzymes, MsIO, MsCPR, MsCYPADH, Ms7DLGT, and Ms7DLH (highlighted in red in **a**), in the middle steps of kratom strictosidine biosynthesis in yeast. Data are presented as mean ± s.d. from three independent biological replicates.

STR-MsCY and STR-CrCY exhibited similar potency in yeast. To further elucidate the specific roles of the three kratom accessory enzymes, we expressed them individually or in combination in the STR-CP5 strain, generating STR-CY3, STR-CY4, STR-CY5, STR-CY6, STR-CY7, and STR-CY8 strains (**Fig. 3b**). Subsequent nepetalactol and tryptamine feeding assays showed that MsCYPADH was essential for 7-deoxyloganic acid production, leading to the accumulation of downstream metabolites (including loganin, secologanin, and strictosidine) in strains expressing MsCYPADH (**Fig. 3b**). Adding MsCPR did not substantially affect intermediate accumulation but markedly increased strictosidine production (a threefold increase to ∼0.06 μM), suggesting that MsCPR enhances metabolic flux through the middle and downstream CYP-catalyzed steps (**Fig. 3c**). In contrast, MsCYB5 had no significant effect. This suggests that either the tested MsCYB5 was not functional under our experimental conditions, or that the step facilitated by MsCYB5 was not rate-limiting in the current design.

Taken together, we identified five additional enzymes, including three pathway enzymes (MsIO, Ms7DLGT, and Ms7DLH) and two accessory enzymes (MsCPR and MsCYPADH). These additions complete the entire kratom strictosidine biosynthetic pathway, with the exception of 8HGO and ISY, which remain unresolved.

### Engineering the yeast endogenous pathway and completing the missing steps toward nepetalactol synthesis

Multiple yeast endogenous enzymes compete with the reactions catalyzed by 8HGO and ISY, diverting geraniol or 8-hydroxylgeraniol toward side products such as citronellol, citronellyl acetate, and 8-hydroxytetrahydrogeraniol, as observed in our previous geraniol feeding assays (**Fig. S5**) and reported in prior studies^47–49^. Therefore, we used CRISPR to delete *OYE2* and *ATF1*, two key endogenous genes competing for geraniol, 8-hydroxylgeraniol, and 8-oxogeranial, from the STR-CP5 strain. The deletion significantly reduced major side-product accumulation upon geraniol feeding in the resulting STR-KO strain (**Fig. 4a**). We then mined the kratom transcriptome to identify more potential enzyme candidates to enable nepetalactol production. NcaMLPLA^21,24^ is a stereoselective cyclase that specifically converts 8-oxocitronellyl enol produced by ISY to the 7S-*cis*-*trans* isomer of nepetalactol. Its expression in yeast and *N. benthamiana* greatly enhanced nepetalactol production compared to spontaneous cyclization. Protein sequence alignment using NcaMLPLA as the query identified a kratom homolog with only ∼40% sequence similarity, which is consistent with reports that MLPLs are largely restricted to the Nepeta genus (**Table S1-S3**). In the meantime, a very recent study reported the discovery of iridoid cyclase (ICYC) in multiple non-Nepeta plants, which also catalyze the cyclization of 8-oxocitronellyl enol to produce nepetalactol stereoisomers^45^. Using the reported *Carapichea ipecacuanha* ICYC as the query, we identified one MsICYC candidate with ∼80% similarity (**Table S1-S3**). While the integration of the putative MLPL with the characterized MsCPR, MsCYPADH, and MsTDC into STR-KO, yielding STR-KI strains, did not yield nepetalactol or other downstream metabolites from either 8-hydroxygeraniol or 8-oxogeranial as expected, the addition of MsICYC led to 103 nM strictosidine when supplied with 8-oxogeranial and tryptamine (**Fig. 4b**). This confirmed the enzymatic activity of MsISY and MsICYC for nepetalactol synthesis. Additionally, genome analysis revealed that the genes encoding MsICYC and the verified MsG8H1 cluster in a 4-kb region on kratom chromosome 11, while MsG8H2, the putative G8H-like candidate that showed no detectable activity in our assays, is on chromosome 10. This organization is consistent with our functional characterization and with the reported conservation of G8H–ICYC biosynthetic gene clusters (BGCs).

**Figure 4.**
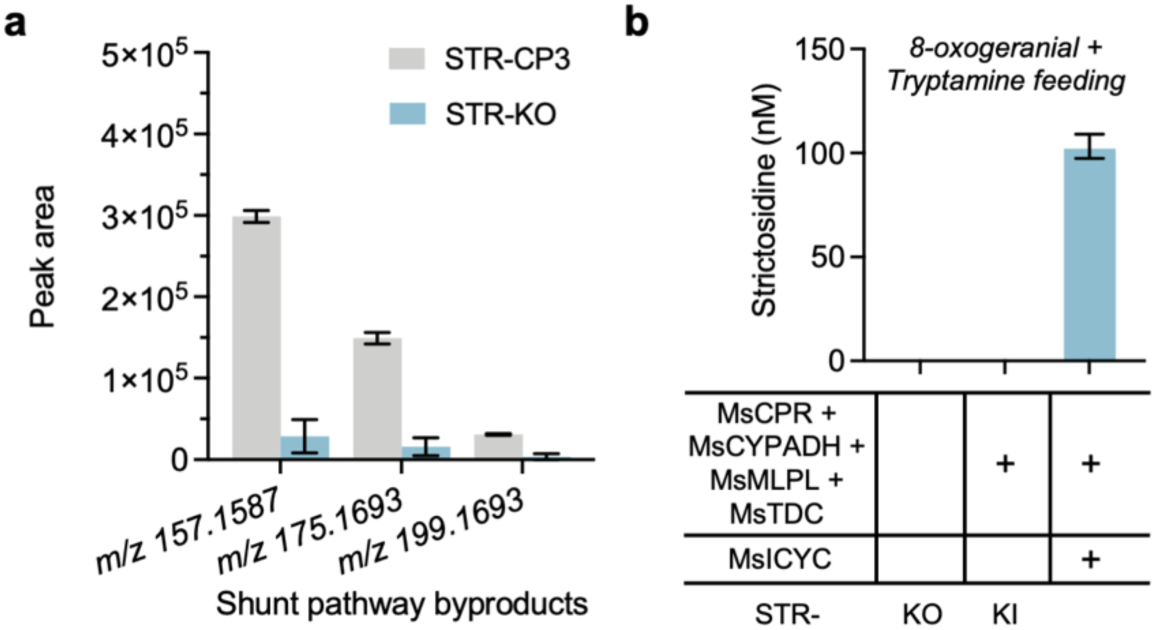
Verification of iridoid synthase and cyclase activity following yeast shunt pathway deletion. **a.** Decreased shunt pathway byproduct production from geraniol and tryptamine after shunt pathway (*OYE2* and *ATF1*) knock-out. STR-KO strains were generated by deleting two shunt pathway genes, *OYE2* and *ATF1*, from STR-CP3 strains, following marker rescue. **b.** Strictosidine production from 8-oxogeranial and tryptamine after MsICYC expression in STR-KI strains from plasmids, validating the function of MsISY and MsICYC. STR-KI strains were generated by integrating MsCPR, MsCYPADH, MsMLPL, and MsTDC genes into STR-KO strains. Data are presented as mean ± s.d. from three independent biological replicates.

After characterizing the activities of MsISY and MsICYC, Ms8HGO remained the only enzyme not yet identified. At the same time, another of our studies revealed that a medium-chain dehydrogenase/reductase from kratom (MsMDR11) can catalyze the interconversions among 8-hydroxygeraniol, 8-oxogeraniol, and 8-oxogeranial *in vitro*. However, this enzyme exhibited primarily reduction activity in yeast, converting 8-oxogeranial to 8-oxogeraniol and 8-hydroxygeraniol (Fig. S7), perhaps related to the highly reducing intracellular environment in yeast.

### Screening of potential transporters via multiplex engineering identified a novel vacuolar importer of secologanin

In parallel, we mined transporters to enhance strictosidine production. Previous studies in *C. roseus* have shown that the strictosidine biosynthetic pathway is spatially organized across multiple cell types and subcellular compartments^9^. Particularly, CrNPF2.6, 2.4, and 2.5 localize to the plasma membrane and mediate the transport of iridoid glucosides, including 7-deoxyloganic acid, loganic acid, loganin, and secologanin^41^. CrNPF2.9 localizes to the tonoplast and exports strictosidine from the vacuole^42^. Using CrNPF2.6 and CrNPF2.9 as queries, we identified two putative kratom homologs, MsNPF2.6 (∼70% identity) and MsNPF2.9 (∼64% identity), from the kratom transcriptome (**Table S1-S3**). We next examined the kratom genome and identified the conserved STR-TDC-MATE BGCs, previously reported in *C. roseus*, *Gelsemium sempervirens*, *Rauvolfia tetraphylla*, and *O. pumila*^33,34,51–53^ (**Fig. S8**). This kratom BGC contained MsSTR, MsTDC, a putative vacuolar MATE transporter that might be involved in the transportation of secologanin (named MsMATE1), as well as two similar putative amino acid transporters homologous to the amino acid transporter AVT1J in *A. thaliana*^54^. Both putative amino acid transporters had low expression levels, and we designated the transporter with relatively higher expression level as MsAVT1J1. Previous studies have suggested that the CrMATE transporter in the cluster may transport secologanin^40,43^. At the same time, a homologous transporter of MsAVT1J1 with 65% identity is also present within the *O. pumila* cluster (**Fig. S8**), both suggesting their relevance to strictosidine synthesis. Using MsMATE1 and MsAVT1J1 as queries, we further identified two additional transporter candidates, MsMATE2 (∼91% identity) and MsAVT1J2 (∼72% identity). Structural modeling with AlphaFold2 revealed that all six kratom transporter candidates contain transmembrane domains^55^ (**Fig. S9**).

We then integrated all six candidate transporter genes, together with the characterized MsCYPADH, MsCPR, and MsTDC, into the STR-KO strain, generating the STR-KT strain. Upon secologanin and tryptamine feeding, the strictosidine titer of STR-KT increased by 38% compared with STR-CP3, rising from 18.42 μM to 25.44 μM (**Fig. 5a**), confirming the transporters’ activities on the transport of secologanin and/or tryptamine, or exporting strictosidine. We attribute this increase in strictosidine production to the introduced transporter set, as the remaining background differences between STR-KT and STR-CP3, namely MsCYPADH, MsCPR, MsTDC expression, and shunt pathway knockout, are not expected to affect the final condensation step.

**Figure 5.**
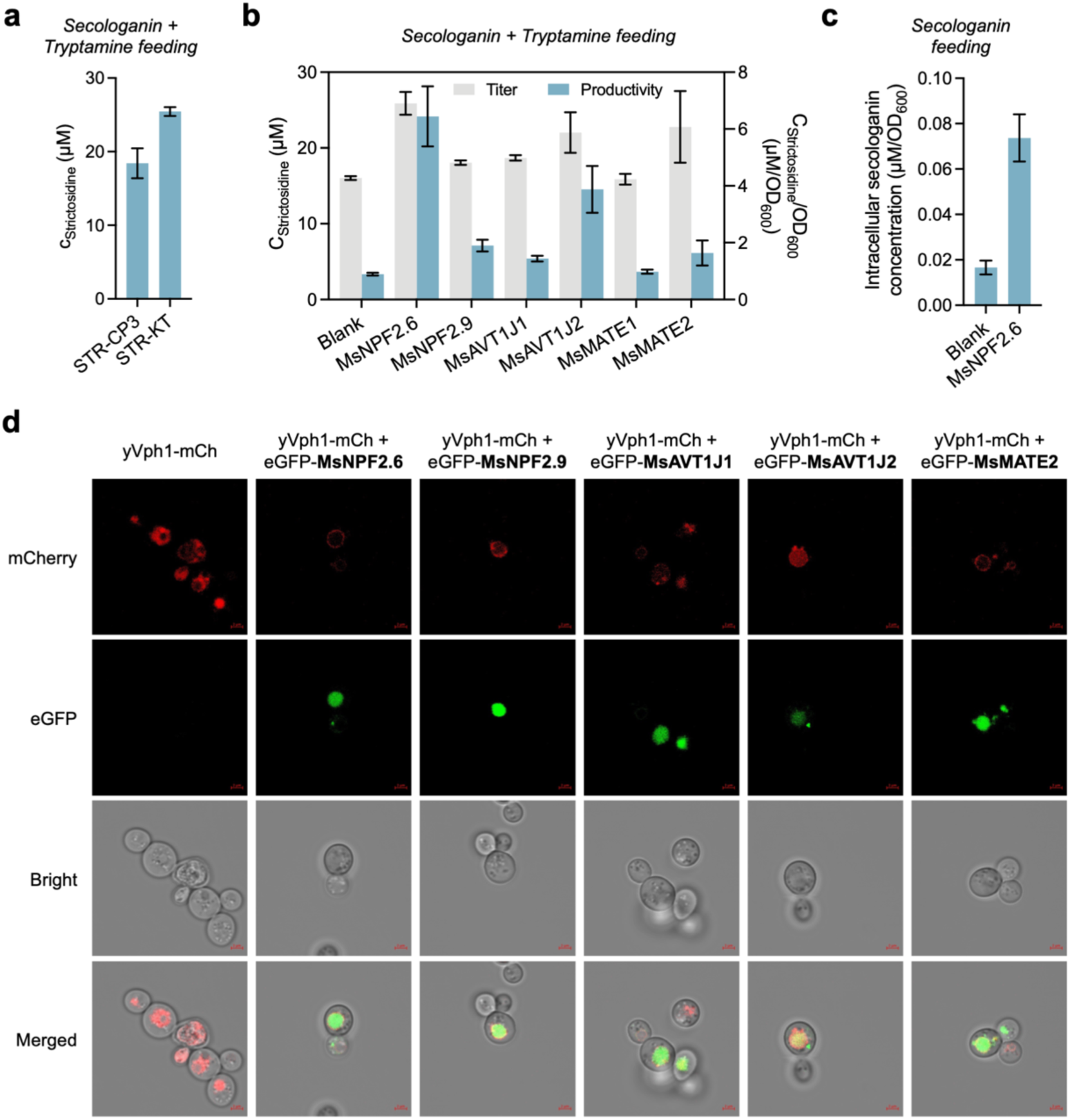
Characterization of potential secologanin and tryptamine transporters. **a.** Results of secologanin and tryptamine feeding assays for STR-KT strains, generated by incorporating MsCPR, MsCYPADH, MsTDC, and six putative transporter genes into the STR-KO strains. **b.** Effects of individual transporters on strictosidine production, assessed by secologanin and tryptamine feeding assays in CEN strains expressing each transporter from plasmids. **c.** Increased intracellular secologanin concentrations in CEN strains expressing MsNPF2.6 as measured by secologanin uptake assays. **d.** Confocal microscopy images of CEN cells co-expressing eGFP-fused transporters and an mCherry-fused vacuolar membrane marker (yVph1-mCh). The red line in the figure represents 2 μM. Data are presented as mean ± s.d. from three independent biological replicates.

To further evaluate the function of the transporters, we expressed the six putative transporters individually in STR-MsCY on plasmids and compared strictosidine production using secologanin and tryptamine feeding assays. All candidates, except MsMATE1, increased strictosidine production and titer at various levels, with MsNPF2.6, MsAVT1J2, and MsMATE2 exerting the highest effects. Compared to the control, expression of MsNPF2.6, MsAVT1J2, and MsMATE2 increased strictosidine titers by 62%, 37%, and 42%, respectively, while MsNPF2.9 and MsAVT1J1 showed modest positive effects on strictosidine titers (**Fig. 5b**). In the meantime, all five transporters led to slower yeast growth, likely due to the compromised membrane integrity due to the high expression of transporters through multi-copy plasmids, as we did not observe a similar level of growth inhibition in STR-KT. Considering the decreased cell density, strictosidine productivity normalized to cell density was further increased. MsNPF2.6, MsAVT1J2, and MsMATE2 increased strictosidine productivity by 620%, 334%, and 84%, and MsNPF2.9 and MsAVT1J1 increased productivity by more than 60%.

We next performed secologanin or tryptamine uptake assays in CEN strains overexpressing the five transporters, respectively. Overexpression of MsNPF2.6 increased intracellular secologanin levels by approximately 3.5-fold (**Fig. 5c**), confirming its function in facilitating secologanin transportation. Confocal microscopic analysis of yeast expressing eGFP-tagged MsNPF2.6 together with an mCherry-tagged yeast vacuolar membrane marker (yVph1-mCh)^56,57^ confirmed that MsNPF2.6 localized to the tonoplast (**Fig. 5d**). Taken together, MsNPF2.6 was designated as a vacuolar importer of secologanin in yeast, unlike CrNPF2.6 that localized to the plant cell membrane in previous studies. Overexpressing MsNPF2.6 increased the concentration of secologanin in the vacuole, consequently enhancing strictosidine production in the vacuole catalyzed by MsSTR. MsAVT1J1, MsAVT1J2, MsNPF2.9, and MsMATE2 also localized to the tonoplast, similar to their *A. thaliana* and *C. roseus* homologs (**Fig. 5d**). The green fluorescence signal was observed both on the tonoplast and in the entire vacuole, supporting our earlier hypothesis that these transporters might be overexpressed and disrupt membrane integrity in yeast. Unlike the vacuolar importer of secologanin CrMATE1, MsMATE2 expression did not alter intracellular secologanin accumulation. We also examined tryptamine import by MsAVT1J1 and MsAVT1J2 but observed no consistent effects. Therefore, the enhancement in strictosidine production could result from enhanced strictosidine export through the tonoplast or the toxicity alleviation and ion homeostasis mediated by these transporters.

### Debottlenecking of kratom strictosidine biosynthetic pathway in yeast with non-kratom genes

We then pursued an alternative strategy to address the bottleneck steps in the elucidated kratom strictosidine biosynthetic pathway in yeast by using non-kratom enzymes. A combination of Vmi8HGO-A, NcaISY, and NcaMLPLA has been shown to address the inefficient production of 8-oxogeranial and nepetalactol in yeast^20,21^. We integrated them into STR-KT strain following the marker rescue, and constructed a new strain STR-KF, which enabled strictosidine biosynthesis from geraniol alone or from combined geraniol and tryptamine feeding in 96-deep-well plate cultures (**Fig. 6a**). In a batch fermentation using shake-flasks (**Fig. 6b**), STR-KF produced strictosidine within the first 48 h, yielding 10 nM with geraniol feeding and 70 nM with combined geraniol and tryptamine feeding. Feeding geraniol together with tryptophan, rather than tryptamine, also resulted in strictosidine production, reaching 20 nM.

**Figure 6.**
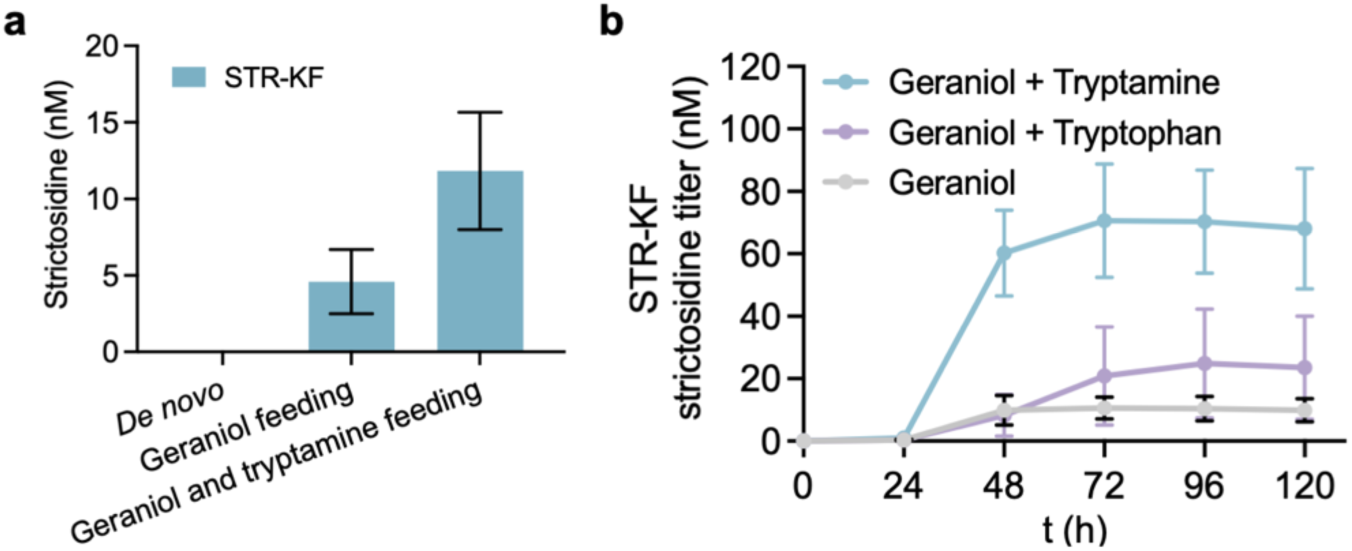
Debottlenecking of kratom strictosidine biosynthetic pathway in yeast with non-kratom genes. **a.** Strictosidine production of STR-KF strains fed with diverse substrates. Integration of non-kratom bottleneck enzyme genes (*Vmi8HGO-A*, *NcaISY*, and *NcaMLPLA*), into STR-KT strains following marker rescue, enabled the strictosidine production from geraniol alone or from geraniol and tryptamine. **b.** Flask-scale fermentation of STR-KF strains fed with diverse substrates. Data are presented as mean ± s.d. from three independent biological replicates.

## Discussion

In this study, we elucidated the kratom strictosidine biosynthetic pathway by identifying at least one functional enzyme of each step in yeast for the first time. Heterologous expression in yeast validated the function of 12 functional enzymes and one novel transporter, including MsGES, MsG8H1, MsISY, MsICYC, MsIO, Ms7DLGT, Ms7DLH, MsLAMT1 and 2, MsSLS, MsCPR, MsCYPADH, and MsNPF2.6. Additionally, MsMDR11 was shown in a separate work to exhibit Ms8HGO activity in vitro, converting 8-hydroxygeraniol to 8-oxogeranial, although in yeast it primarily catalyzed the reverse, reductive reactions. Leveraging these enzymes, we were able to produce 103 nM strictosidine from 8-oxogeranial and tryptamine. These enzymes provide novel building blocks for future MIA pathway reconstruction and engineering and enrich current database of MIA enzymes. We have also addressed the bottleneck step by using genes from other plants, resulting in a strictosidine titer of 70 nM from geraniol and tryptamine. This discovery has showcased the biochemical conserveness of early steps in MIA metabolism across different MIA-producing plants, since all edit later kratom enzymes share high functional and sequence identity (over 74%) with those identified in *C. roseus*. In contrast, elucidated enzymes downstream of strictosidine, even those catalyzing the same type of enzymatic transformations, exhibited significantly lower sequence identity. For example, MsSGD shares only 55% sequence identity with CrSGD, and enzymes catalyzing heteroyohimbine synthesis, such as tetrahydroalstonine synthase and heteroyohimbine synthase, exhibit only 52% and 64% identity^11^. This reflects the MIA diversification between the Ruberaceae and the Apocynaceae families.

Previous studies in *C. roseus* have shown that the strictosidine biosynthetic pathway is spatially organized across multiple cell types and subcellular compartments^9^. Particularly, the intracellular transport of intermediates across multiple organelles is vital to the biosynthetic efficiency and cell robustness. Enzymes catalyzing the syntheses of secologanin and tryptamine are localized either in the cytoplasm or on the endoplasmic reticulum (ER) membrane facing the cytoplasm. In contrast, the last step condensing secologanin and tryptamine takes place in the vacuole^9,58^. Therefore, vacuolar transporters that can import secologanin and tryptamine and export strictosidine are particularly important in both natural and engineered strictosidine production. The functional validation of MsNPF2.6 as a secologanin transporter provides experimental support for the long-proposed role of NPF2.6 in plant strictosidine biosynthesis.

Elucidation and optimization of the kratom strictosidine pathway were markedly accelerated by multiplex pathway engineering, which enables the simultaneous genomic assembly and expression of multiple candidate genes in yeast. This strategy enables functional validation of the entire pathway in parallel rather than sequentially, substantially reducing the time and experimental burden associated with conventional step-by-step characterization. Multiplex methods such as HI-CRISPR^59^, COMPASS^60^, and CasEMBLR^61^ have demonstrated the feasibility of reconstructing known pathways in yeast. We further demonstrated this strategy’s potential for elucidating unknown pathways. By testing multiple enzymes that catalyze sequential reactions as a functional module, this approach enables rapid validation of productive combinations while efficiently eliminating incorrect candidates. The ability to reject nonfunctional hypotheses early, rather than investing in prolonged stepwise characterization, is particularly valuable for complex plant pathways with large gene families and ambiguous annotations. As such, multiplex pathway engineering represents a powerful and underutilized framework for accelerating both the discovery and optimization of plant natural product biosynthetic pathways in yeast.

## Methods

### Chemicals and reagents

Yeast nitrogen base (YNB) and amino acid mixtures were purchased from Sunrise Science Products. Ammonium sulfate, dithiothreitol (DTT), tris(2-carboxyethyl) phosphine hydrochloride (TCEP), secologanin, and tryptamine were purchased from Sigma-Aldrich.

Dextrose, yeast extract (YE), peptone, Luria-Bertani (LB) broth, agar, acetonitrile, and formic acid were purchased from Thermo Fisher Scientific. Geraniol was purchased from Alfa Aesar. 8-Hydroxygeraniol and cis/trans-nepetalactol were purchased from Santa Cruz Biotechnology. 8-Oxogeranial was purchased from Toronto Research Chemicals. Loganic acid and loganin were purchased from Cayman Chemical. All other chemicals, including antibiotics, were purchased from VWR International.

### Kratom gene mining

The *C. roseus*, Nepeta, and *C. ipecacuanha* protein sequences were used to identify counterpart sequences in the kratom transcriptome (ref). The top hits were selected for gene integration and biochemical characterization. All sequences used for gene mining in kratom are listed in **Supplementary Table S1**. Candidate gene sequences and alignment results are listed in **Supplementary Table S2** and **S3.**

### Kratom leaf RNA extraction and cDNA preparation

Kratom leaf RNA purification and cDNA preparation were performed as previously described^11^. Briefly, mature (six-week-old) and young (two-week-old) leaves were harvested, flash-frozen in liquid nitrogen, and ground into a fine powder. Total RNA was extracted using the RNeasy Plant Mini Kit (QIAGEN) according to the manufacturer’s protocol. cDNA was synthesized from 3 µg of leaf RNA using the RNA to cDNA EcoDry Premix (Takara Bio), following the manufacturer’s instructions. The resulting cDNA was purified with the DNA Clean & Concentrator Kit (Zymo Research), yielding approximately 180 ng of cDNA per sample.

### Plasmid construction

Gene sequences used in this study are listed in **Supplementary Table S2** and **S4.** Plasmids used in this study are listed in **Supplementary Table S5**. For gene expression cassette assembly, genes and pE holding plasmid backbones, containing specific promoter-terminator pairs, were amplified using Q5 High-Fidelity DNA Polymerase (New England Biolabs) with primers from Life Technologies, purified by agarose gel extraction with the Zymoclean Gel DNA Recovery Kit (Zymo Research), and assembled by Gibson Assembly^62^ with the NEBuilder HiFi Assembly Master Mix (New England Biolabs). Kratom leaf cDNA, Yeast Strain 4 genomic DNA^39^, synthesized codon-optimized gene fragments (Twist Bioscience), and previously constructed holding plasmids^44^ were used as templates for putative kratom genes, *C. roseus* CYP accessory enzymes (*CrCYR*, *CrCYB5*, and *CrCYPADH*), other non-kratom genes (*ATR1*, *Vmi8HGO-A*, *NcaISY*, and *NcaMPLPA*), and holding plasmid backbones, respectively. For yeast gene expression plasmid construction, gene expression cassettes on pE vectors were transferred into multicopy yeast expression plasmids via Gateway LR recombination using the Gateway LR Clonase II Enzyme mix (Life Technologies). For Cas9 plasmid construction targeting yeast genome, the crRNAs, direct repeats, and spacers were synthesized by Life Technologies and assembled into pCRCT plasmid containing a Cas9 expression cassette with the Golden Gate method by NEBridge Golden Gate Assembly Kit (New England Biolabs).

*E. coli* competent cells were prepared with the Mix & Go! *E. coli* Transformation Kit (Zymo Research) and used for the plasmid construction. Plasmids were purified using the ZR Plasmid Miniprep kit (Zymo Research), and the sequences were confirmed by Sanger Sequencing (Biotechnology Resource Center, Cornell University) or Nanopore sequencing (Plasmidsaurus).

### Microbes and culture conditions

Strains used in this study are listed in Supplementary Table S6. *E. coli* Top 10, ccdB resistant *E. coli*, Yeast Strain 4^39^, and yeast strains derived from CEN.PK2-1D (CEN) were used in this work. *E. coli* strains were cultivated in LB media at 37 °C with 50 µg/mL kanamycin or 100 µg/mL carbenicillin, as appropriate. Unless specifically mentioned, yeast strains were grown at 30 °C in YPD media (1% yeast extract, 2% peptone, 2% dextrose) or in synthetic dropout (SD) media (0.17% yeast nitrogen base, 0.5% ammonium sulfate, 2% dextrose, and appropriate amino acid dropout mixture). For solid media, 2.5% agar was added.

### Genomic integration of biosynthetic pathways and shunt pathway deletions in yeast

The MULTI-SCULPT^44^ method was used for multiplex genomic integration of biosynthetic pathways into the yeast genome, as previously described. Briefly, gene expression cassettes, selective marker gene cassettes, and 500-bp upstream and downstream genomic homologies of each integration site, each harboring 25-bp overlaps between adjacent fragments, were first amplified from holding plasmids or yeast genomic DNA, using Q5 High-Fidelity DNA Polymerase (New England Biolabs) with primers synthesized by Life Technologies. PCR products were then purified by agarose gel extraction with the Zymoclean Gel DNA Recovery Kit (Zymo Research), quantified with a Nanodrop spectrophotometer (Thermo Fisher Scientific), mixed in equimolar ratios (500 fmol per insert) with 1000 ng of the corresponding Cas9 plasmid, and concentrated to 5 μL by Savant SpeedVac DNA130 (Thermo Fisher). The mixture was electroporated into 50 μL of yeast electrocompetent cells in a 2 mm cuvette using a Gene Pulser Xcell Total System electroporator (Bio-Rad) at 540 V, 25 μF, and infinite resistance. Transformants were immediately recovered in 1 mL of 2×SD-URA medium at 30°C and 400 rpm overnight, then centrifuged and plated onto the appropriate SD plates for 4 days. Correctly assembled yeast strains were verified through colony PCR. For subsequent rounds of multiplex integration, the engineered parental yeast strains were transformed with Cre recombinase plasmids to excise the integrated selective marker cassettes, cultured in SC medium containing 5-fluoroorotic acid and 2-amino-5-fluorobenzoic acid to remove the Cre recombinase and Cas9 plasmids, and used for the next round of multiplex integration following the procedure described aboveShunt pathway deletions were performed following the same MULTI-SCULPT and marker excision protocols. Cas9 target sites and crRNA sequences are listed in **Supplementary Table S7**. Standard pathway integration at a single genomic locus was carried out similarly but without Cas9 plasmids.

### Plasmid-based gene expression in yeast

Yeast gene expression plasmids (500 ng each) were transformed into chemically competent yeast cells prepared using the Frozen-EZ Yeast Transformation II Kit (Zymo research) according to the manufacturer’s protocol and plated onto the appropriate SD plates.

### Yeast fermentation and sample preparation

Yeast colonies from the plates were picked and cultured in YPD, SC, or corresponding SD medium at 30 °C and 400 rpm for 48 h as seed cultures. Seed cultures were then back-diluted 10-fold into fresh corresponding medium supplemented with substrates (1 mM for tryptamine and 0.25 mM for other substrates, unless otherwise specified) and 25 mM NaOH and incubated for 72 h at 30 °C and 400 rpm (500 μL total volume for fermentations in 96-well plates and 25 mL for fermentations in flasks). For fermentations in 96-well plates, cultures were centrifuged at 14,000 rpm for 15 min, and the supernatants were used for metabolite analysis. For flask-scale fermentations, 1 mL of culture was adjusted to pH 10 with NaOH, mixed with three volumes of ethyl acetate, and vortexed. The ethyl acetate layer was collected and evaporated by Savant SpeedVac DNA130 (Thermo Fisher), and the residue was resuspended in 75 μL of methanol for metabolite analysis. Fermentations were performed in triplicates.

### Metabolite analysis of yeast fermentation products

Metabolites in fermentation products were identified and quantified using HPLC/Q-TOF (Agilent 1260 Infinity II/Agilent G6545B) in MS mode with positive ionization. For analysis, 5 μL of sample was injected and separated in the Agilent ZORBAX RRHD Eclipse Plus C18 column (2.1 × 50 mm, 1.8 μm) with mobile phase A (water containing 0.1% formic acid) and B (acetonitrile containing 0.1% formic acid). The gradient program was set as follows: 0-1 min, 95% A; 1-11 min, 95%-5% A; 11-13 min, 5% A; 13-14 min, 5%-95% A; and 14-16 min, 95% A, at 40°C and a flow rate of 0.4 mL/min. The m/z value was used to extract the ion chromatogram (EIC, ± 10 ppm) for compound identification and quantification. For metabolites with available chemical standards, concentrations were determined by comparing integrated peak areas to standard curves. For putative metabolites, EIC peak areas were used for relative quantification.

### Secologanin and tryptamine uptake assays

Yeast expression plasmids containing transporters or blank control were transformed individually into CEN, and the resulting strains were cultivated in the appropriate SD media at 30°C and 400 rpm for 48 hours. Cell density was measured, and the cultures were then centrifuged and concentrated to an OD_600_ of 10 in 0.5 mL of fresh appropriate SD media (100 mM Tris, pH=7.5) containing either 0.1 mM secologanin or 1 mM tryptamine. The cells were incubated in 96-well deep-well plates at 30°C and 400 rpm for 24 hours. The cell density was measured, and cells were then harvested, washed twice with ice-cold water, and resuspended in 100 μL of water. The resuspended cultures were lysed in microtubes containing 425–600 μm glass beads (Sigma-Aldrich) using a BeadBug 6 microtube homogenizer (Benchmark Scientific) for twelve 30 s bursts, with 30 s rests between cycles. After lysis, 100 μL of water was added. The cell lysates were then centrifuged at 15,000 rpm for 15 min, and the supernatants were used for HPLC/Q-ToF analysis to quantify secologanin and tryptamine concentrations, which were normalized to OD_600_ as a proxy for intracellular concentration. Uptake assays were performed in triplicates.

### Characterization of subcellular localization of transporters in yeast

Plasmids expressing the mCherry-tagged yeast vacuolar membrane marker (yVph1-mCherry) and eGFP-tagged transporters were co-transformed into CEN. Resulting yeast strains were cultivated at 30°C, 400 rpm in appropriate SD media for 24 hours and imaged using a Zeiss LSM 710 Confocal Microscope (AxioObserver, with objective Plan-Apochromat 63X/1.40 Oil DIC M27).

## Acknowledgements

We thank the Flow Cytometry Facility (RRID: SCR_021740) and the Imaging Facility (RRID: SCR_021741) of the Biotechnology Resource Center of Cornell Institute of Biotechnology. We thank J. Han and C. Liu for their helpful suggestions and discussions. This work was supported by National Institutes of Health grant R01AT012633, National Science Foundation grant MCB-2338009, DBI-2019674, and IOS-2220733, Schwartz Research Fund Award, Cornell Engineering SPROUT Award, and Cornell CALS Moonshot Seed Grant.

## Author contributions

YW, DH, FG, and SL conceived and designed the research. YW, DH, and FG performed the experiments and analyzed the data. YW, DH, FG, and SL wrote the paper. All authors read and approved the final manuscript.

